# Emergent coexistence in multispecies microbial communities

**DOI:** 10.1101/2022.05.20.492860

**Authors:** Chang-Yu Chang, Djordje Bajic, Jean Vila, Sylvie Estrela, Alvaro Sanchez

**Affiliations:** Department of Ecology & Evolutionary Biology, Yale University, New Haven, CT, USA; Microbial Sciences Institute, Yale University, New Haven, CT, USA

## Abstract

Microbial communities are highly diverse, and understanding the factors that promote and modulate this diversity is a major area of research in microbial ecology. Recent work has proposed a reductionist perspective to microbial coexistence, where pairwise coexistence between strains in isolation is required for their coexistence in a more complex community. In this view, species exclusion in pairwise co-culture would preclude their coexistence in a more complex community too. An alternative view is that coexistence is a more complex property of the entire community, requiring the presence of additional community members. If this view were correct, competitive exclusion in pairwise co-culture would not necessarily preclude species coexistence in more complex community contexts. Empirically testing these alternative hypotheses is complicated by the intractably high microbial diversity of most natural communities, and the challenges of reconstituting every pair of coexisting species under the exact same habitat where their community of origin was assembled. To address this challenge, we have experimentally reconstituted all possible pairwise co-cultures between stably coexisting species from 13 different, low-diversity microbial enrichment communities, which had previously been assembled in a well-controlled synthetic habitat. We find that, when isolated from the rest of their community members, most species pairs fail to coexist. This result highlights the importance of community context for microbial coexistence and indicates that pairwise exclusion may not reflect the ability of species to coexist in more complex, multispecies ecosystems.

## INTRODUCTION

Explaining species coexistence and the bewildering diversity of ecological communities has been one of the major goals of Ecology^1^. Historically, research on the mechanisms that make coexistence possible has focused on pairs of competing species^2^. This work has been instrumental to develop our current understanding of the fundamental mechanisms that modulate biodiversity, and it occupies a central place in ecological theory. Yet, species are found in nature forming complex ecological communities. It is thus plausible that their coexistence may be mediated by interactions that require the presence of more than two species, including higher-than-pairwise interactions, but also complex networks of pairwise effects^3–7^.

Is species coexistence primarily a reductionist pairwise phenomenon, or is it an emergent property of the whole community (Fig. 1)? In other words, does coexistence within a complex community require that two microbial species also coexist in isolation? To unambiguously answer this question, one would have to break down a stable multispecies community into its simplest units (species pairs), and determine whether all of those pairs coexist too. If we found that all (or, more generously, most) of them did, this would lend support to the reductionist hypothesis. If, on the other hand, we found that few of those species coexist as a pair when removed from the ecological context provided by the other community members^8^, we would have to conclude that multispecies coexistence is an emergent, systems-level property of the community as a whole. An extreme scenario is the maintenance of biodiversity through nontransitive (non-hierarchical) competition, which may allow multiple species to coexist even when none of them do as a pair in isolation^4,5,9–12^.

**Figure 1.**
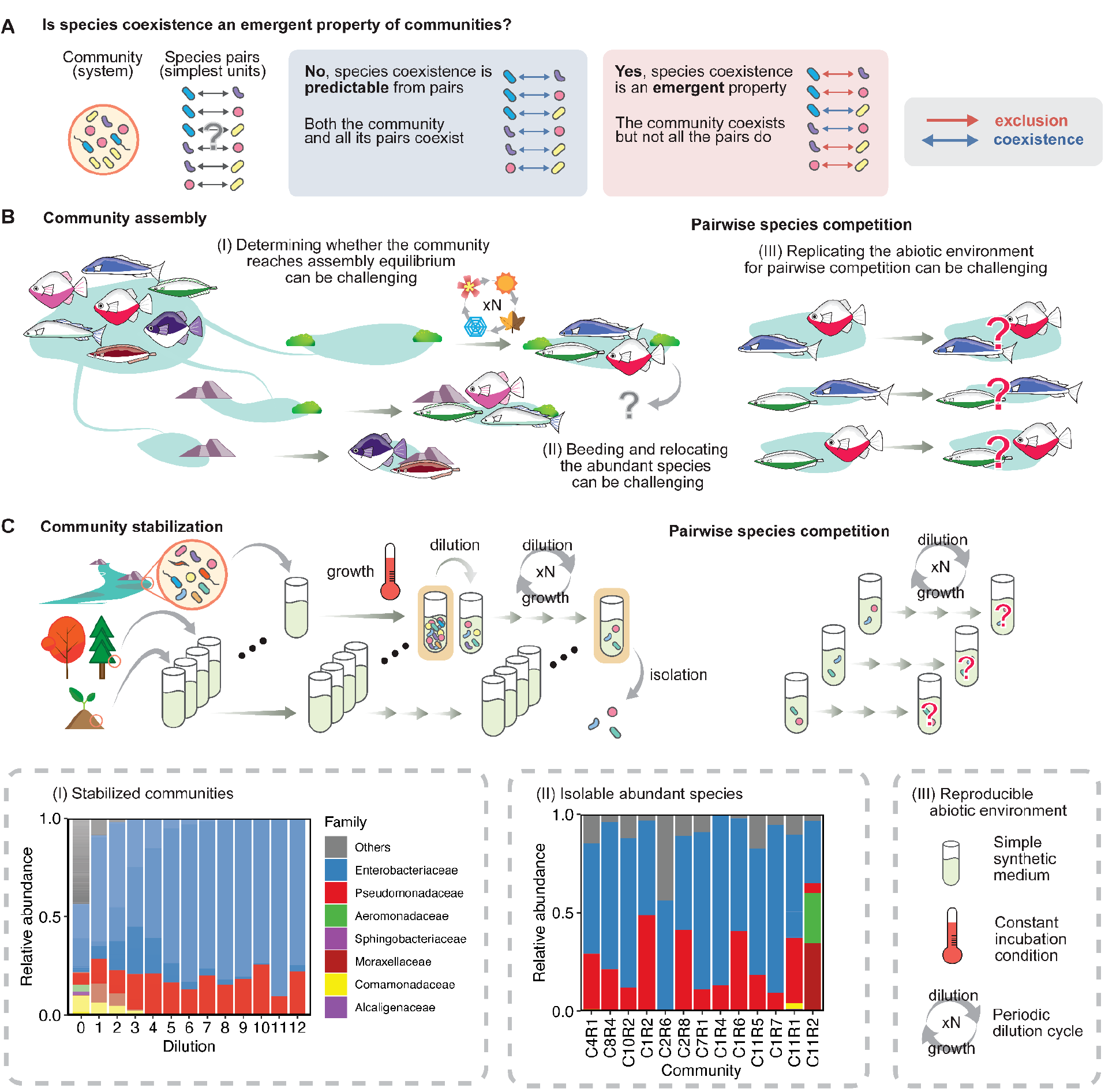
Empirically testing whether species coexistence is an emergent property of communities. (**A**) An emergent property occurs when it emerges at the system level (community) but is not observed at the level of its elemental units (species pairs). If all species isolated from a community coexist with one another in pairs, the coexistence at the community level is predictable from the knowledge of pairs, therefore can be reducible to simple additivity of all pairs. Whereas when some pairs do not coexist, species coexistence is an emergent property of the diverse communities that requires the concomitant presence of more than two species. (**B**) To empirically test this question in a set of natural ecosystems, one needs to be able to breed the individual species from each community, and reconstitute the competing species pairs. However, these procedures can be logistically challenging regarding the community assembly stage, species isolation, and environment reconstruction. (**C**) To address the challenges described above, we have developed an empirical system including hundreds of microbial communities assembled in a reproducible abiotic environment, allowing a direct test of emergent species coexistence. These communities had been stabilized in a simple synthetic medium with one limiting nutrient at a constant incubation condition (Methods). The analysis of the community composition of a representative community (C8R4) after every dilution cycle shows stabilization. Color shades represent different ESVs. We chose 13 representative communities that cover a range of strain diversity (N=3-12) for examining the pairwise competition outcomes and were able to match the isolated strains to the community ESVs which explains an average of 89.5% of relative abundance (inset II; Methods).

Resolving this question is particularly urgent in microbial ecology, given the enormous and still largely unexplained diversity of microbial ecosystems^13–20^. The answer is likely going to be system-specific. Recent empirical research has provided support to the reductionist paradigm, finding that the composition of synthetic bacterial communities can be predicted from a simple pairwise assembly rule: the species that coexist together in a complex community are those that coexist with one another as pairs in isolation^21,22^.

However, synthetic bacterial communities have also exemplified nontransitive loops (i.e., rock-paper-scissors) where a trio coexists despite all pairwise co-cultures ending up in exclusion^23,24^. Testing these two scenarios in natural communities is generally not feasible, given their diversity: even if we managed to isolate most community members from a natural habitat, the number of replicate natural environments we would need in order to incubate every single pair would scale quadratically with the community richness.

Given the technical, financial, and even ethical difficulty of doing such an experiment in environmental or medical settings, previous studies took an inference-based approach where they estimated the pairwise interaction coefficients by fitting models of population dynamics to the temporal behaviors of the community and/or a subset of sub-communities^25,26^. From these models, one may infer the outcome of pairwise co-cultures. Despite its mathematical elegance, generating predictions from this approach requires a large collection of data and may be prone to artifacts and failed predictions, as the models generally assume that high-order interactions are absent and impose specific functional forms on the density-dependence of pairwise interactions. As an alternative approach, recent studies have attempted to directly determine pairwise coexistence by reconstituting species pairs from natural communities in well-controlled laboratory environments^21,27,28^. A limitation, however, was that only a very small subset of all of the coexisting species in those natural communities was included in the study, typically those which could grow on their own under laboratory conditions. Perhaps most importantly, the pairs were reconstituted in a laboratory environment that is very different from the original environment experienced by the community where those species pairs were coexisting. Thus, while pairwise assembly rules were generally successful at predicting the final composition of small (N<5) synthetic consortia in novel environments, they did not permit one to determine whether or not multispecies coexistence in the communities these isolates came from was a pairwise phenomenon.

In past work, we have assembled a collection of hundreds of microbial enrichment communities that stably coexist in well-controlled synthetic environments containing one single supplied limiting nutrient^15,29–32^. These communities were formed in a manner that is very similar to the “random zoo” model in theoretical ecology^33^: we added a large number of species to a new habitat (which none of them had encountered before) without giving them much time to adapt and allowed these species to spontaneously assemble into a new stable community (Fig. 1C). These newly assembled communities reach compositional stability after ~50-80 generations of serial passaging. Once stabilized, they have a modest diversity (N<25, generally^29,30,32,34^), and most of the species can be isolated without much work, including those that cannot grow on their own^29,30^. Because the communities were assembled in liquid, synthetic media under laboratory conditions that can be easily and faithfully recreated in high throughput, it is possible to break each community apart and reconstitute every pairwise species combination in exactly the same habitat where the multispecies community from which they came from was assembled in the first place. This is critically important, as it allows us to evaluate whether a pair coexists in isolation under identical abiotic conditions where they coexist as a part of a multispecies community. All of these advantages overcome the limitations of past laboratory studies, providing a middle ground between natural and purely synthetic communities^31^.

Motivated by this realization, we set out to test whether coexistence is a pairwise or an emergent phenomenon in this empirical system. To that end, we empirically characterized the competition outcomes in all 186 within-community pairwise combinations among 68 microbial species from 13 representative multispecies enrichment communities, comprising ~90% of all the Exact Sequence Variants (ESVs) in those communities^30^. As shown below, we find that most species pairs fail to coexist in isolation under the same ecological conditions where they do coexist as members of the same multispecies community. Among those pairs that do coexist, stable rather than neutral coexistence is the norm. Among non-coexisting pairs, competitive exclusion is strongly hierarchical, a point that is consistent with previous findings^28^.

## RESULTS

### Multi-replicated enrichment microbial communities are an ideal model system to test whether multispecies coexistence is a pairwise or emergent property

Over the past few years, we have gathered a collection of hundreds of self-assembled communities in well-controlled synthetic habitats^29^. Starting from natural bacterial communities that were serially passaged every 48 hr in glucose minimal medium (Methods), these enrichment communities adopted stable compositions containing N<25 Exact Sequence Variants (ESVs) after ~50 generations. The simple synthetic medium and incubation conditions make it straightforward to reconstruct the abiotic environment, allowing us to characterize the pairwise competition outcomes under the exact abiotic conditions where the original community had been assembled.

To empirically test whether multispecies coexistence is a pairwise phenomenon in our system, we chose 13 representative communities containing 3-12 isolated strains. We were able to isolate 68 bacterial strains whose full-length 16S rDNA matched the majority of the relative abundances obtained from the exact sequencing variants (89.5% of relative abundance). Pairwise competition experiments were then carried out by mixing pairs of the isolates’ inocula with standardized optical density (OD=0.1 at 620 nm) at three sets of given volumes (the cell proportion then approximately 5:95, 50:50, and 95:5), and passaging them for eight growth-dilution cycles in the same glucose minimal medium at the same 30°C (Fig. 2A). During each growth cycle, the cells were incubated for 48 hr, after which the resulting culture was diluted by 1/125 into fresh media as we did in the original community assembly experiment. At the end of the last dilution cycle, we measured the composition of pairwise communities by plating the cocultures on petri dishes and counting the colonies. To avoid human bias in colony morphology identification, we adopted an automated image processing pipeline (Fig. S1) combined with a machine-learning approach for classification on a total of 186×3=558 coculture images using 40 colony morphology features (Fig. S2-3; Table S1; Supplement Materials PDFs). Among 186 competing pairs of strains, we removed six pairs because these cocultures did not grow and has no colony (Methods). We also removed another nine pairs where the trained model performs poorly on the validation datasets (accuracy score < 0.9; Methods; Fig. S4), resulting in N=171 pairs in our analysis. The machine result also corresponds to the human result for both the total colony count on a plate (R^2^=0.88) and the relative frequency of colony morphotypes (R^2^=0.85) (Fig. S5).

**Figure 2.**
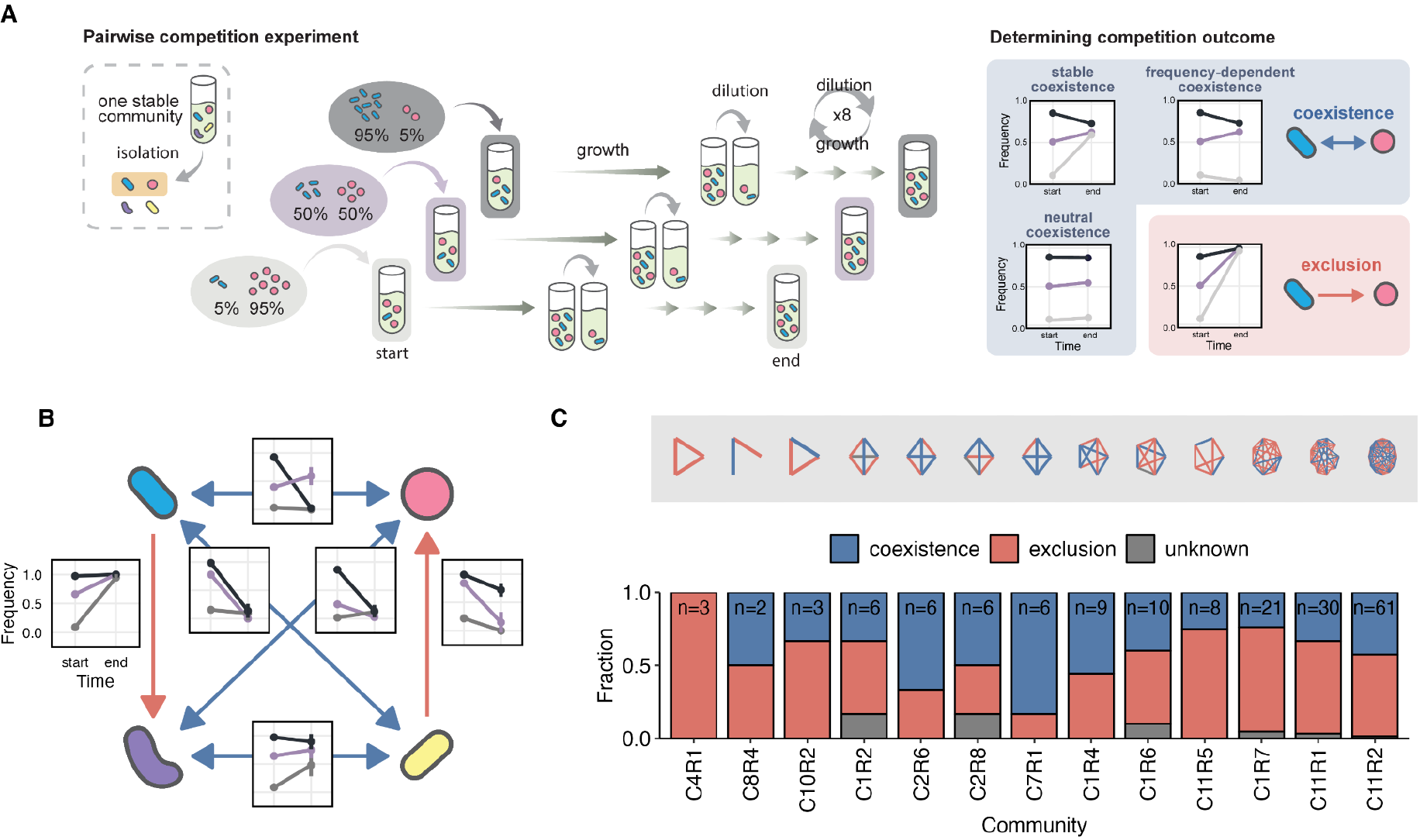
Multispecies coexistence is an emergent property as most species pairs fail to coexist on their own. **(A)** We show a hypothetical example of one species pair from the pairwise competition experiment. For a pair of strains, the competition outcome was determined by competing them in three varied initial frequencies to test whether they are capable of invading each other from rare. The competition outcome was determined by a mutual invasion criterion that has been used in previous studies^21,27^: a pair stably coexists when one strain can invade the other from rare and vice versa; whereas if one strain always excludes another regardless of its initial frequencies, this pair was deemed as a competitive exclusion pair. We also included neutrality as coexistence in a broad sense. (**B**) Analysis of the pairwise competition outcomes of a representative community (from community C2R6) reveals a combination of both pairwise exclusion and pairwise coexistence among the four constituent taxa. Each inset line chart shows the frequency changes between the start (day 0) and the end (day 8) of the competition experiments in a pair. The values for each point and its error bar are the mean and standard deviation of 1000 bootstraps (Methods). The color intensity in the insets represents the set of initial frequencies used in the pairwise competition: 95% (darkest purple), 50% (medium purple), to 5% (lightest purple). The pairwise competition outcomes of all 13 enrichment communities are shown as the network in (**C**), ordered by the number of strains in each community from the smallest (3 taxa) to the largest (12 taxa). The sample size in each bar shows the number of pairs per network. Some networks have missing pairs because these pairs either do not have any colonies in cocultures or have low classification model accuracy.

In general, a pair is identified as “frequency-dependent coexistence” if there is a sign of negative frequency-dependent coexistence, meaning that both species could invade each other when rare^21,35^. We also consider neutral coexistence if the relative frequency does not change significantly over the course of experiments. By contrast, competitive exclusion is deemed to be the outcome when one of the species drops in abundance below the detection limit of our experiments regardless of its initial frequency.

### Species coexistence is an emergent property in our enrichment microbial communities

To recapitulate the experiment described in Fig. 1C, we reconstituted every possible pair of species from each of the 13 communities, in the exact same environment where the community had been assembled (Fig. 2A). Thus, all of these pairs consisted of two species that coexist together in the context of a more complex community. After characterizing the outcome of every one of these 186 pairwise co-cultures and removing 15 inapplicable pairs because of the lack of colonies or low model accuracy (Methods), we found that competitive exclusion was the most common outcome 97/171=56.7%, and only 68/171=39.8% of the species pairs were able to coexist (stably or neutrally) in pairwise co-culture when isolated from the rest of the species in the community. Note that we have six pairs (6/171=3.5%) that exhibit strange fitness functions and were left unknown. All of the 13 communities contained at least one pair (but generally many more) that did not coexist (Fig. 2B) and the fraction of pairs exhibiting exclusion was remarkably consistent and comparable for all of the 13 enrichment communities, regardless of their richness (Fig. 2C). To avoid that pairwise competition outcomes may be dictated by pairs of rare strains (relative abundance < 5%), we filtered for pairs containing only the highly abundant strains and repeated the same analysis. The result holds quantitatively: exclusion occurs at 47/79=59.5%, coexistence at 29/79=36.7%, and unknown at 3/79=3.8% (Fig. S6).

Our results indicate that the simple pairwise assembly rule proposed in ref. ^21^ would have failed when applied to our consortia, as it would have predicted a much less diverse set of communities composed only of those taxa that coexist as an isolated pair. This result suggests that, at least under our experimental conditions, multispecies coexistence is a complex, emergent property of the entire community that cannot be trivially reduced to the outcome of pairwise co-cultures in isolation. This suggests that the outcome of competition experiments between pairs of isolates in the absence of additional community members has a more limited value than previously thought, at least when we seek to predict community composition.

Among the coexisting pairs, the majority were found to coexist stably (Methods), as evidenced by negative frequency-dependent selection: both species were able to invade each other when rare (“stable coexistence” 26/171=15.2% or “frequency-dependent coexistence” 3/171=1.8%) (Fig. 3). A considerable number of pairs coexisted at a low abundance (coexistence at 5% or 95%, each is 17/171=9.9% and 18/171=10.5%) but we could not detect negative frequency dependence. Interestingly, neutral coexistence among isolated pairs also occurred but was rare, with four pairs of species exhibiting dynamics that did not rule out neutrality (Methods).

**Figure 3.**
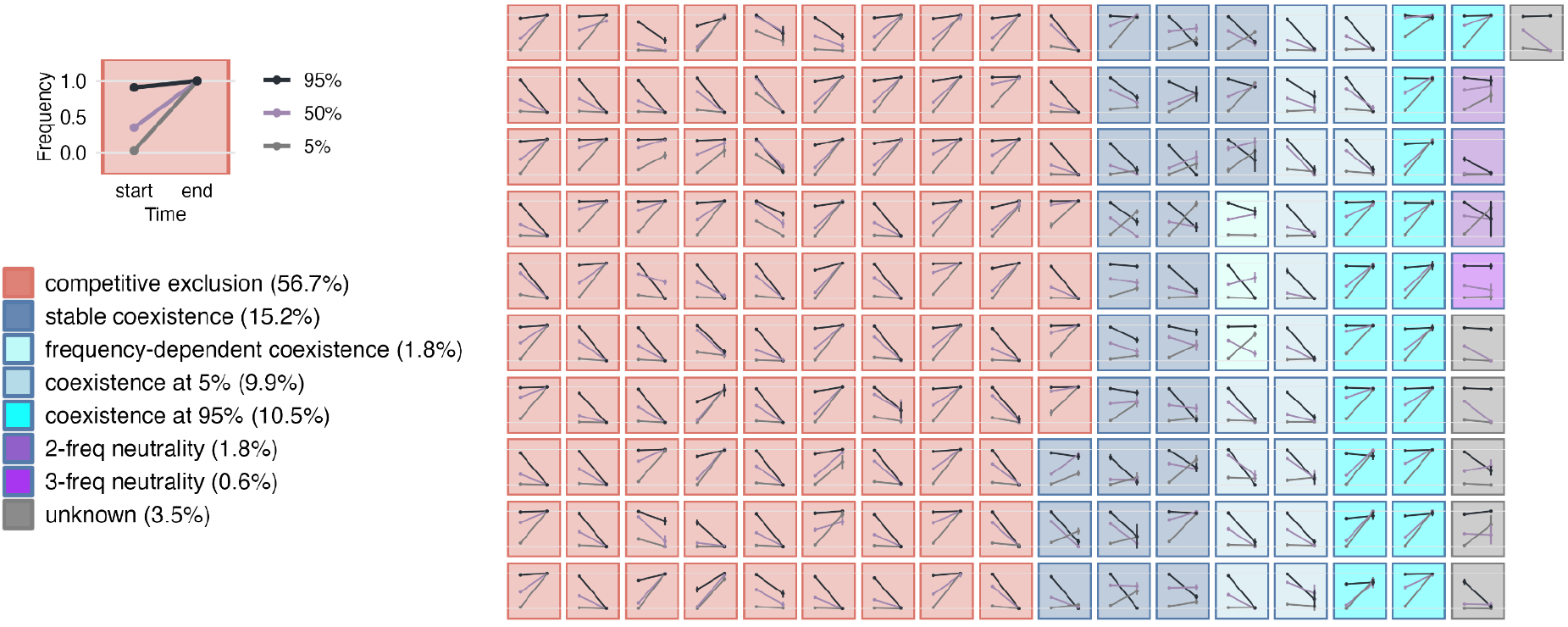
Species pairs from coexisting communities exhibit exclusion, frequency-dependent coexistence, and neutral coexistence. Competition outcomes of a total of 171 pairs across the 13 enrichment communities. The values for each point and its error bar are the mean and standard deviation of 1000 bootstrapping samples (Methods). Each of the inset grids indicates the dynamic change from the start (day 0) to the end (day 8). The background color represents the identified competition outcomes whereas the line color indicates the treatments of three initial frequencies.

### Competitive exclusion among species pairs in isolation is transitive and strongly hierarchical

An extreme case of emergent coexistence may occur when coexistence networks are nontransitive. An example is the “rock-paper-scissors” scenario, where species A outcompetes species B, species B outcompetes species C, and species C outcompetes species A. To determine whether this scenario might play a role in stabilizing coexistence in our communities, we ordered all strains in each community by their competitive ranking, which is calculated as how many times an isolate won in a pairwise competition minus how many times it lost, divided by the number of games it has played (Fig. 4A; Methods). Regardless of the community size, we observed a consistent hierarchical structure in all 13 communities when these strains are ordered in their rankings (Fig. 4B). To quantify the hierarchies of our communities, we introduce a simple metric (*h*) which captures whether the direction of pairwise exclusion (from winner to loser) conforms to the strains’ competitive ranking:

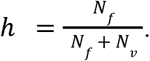

**Figure 4.**
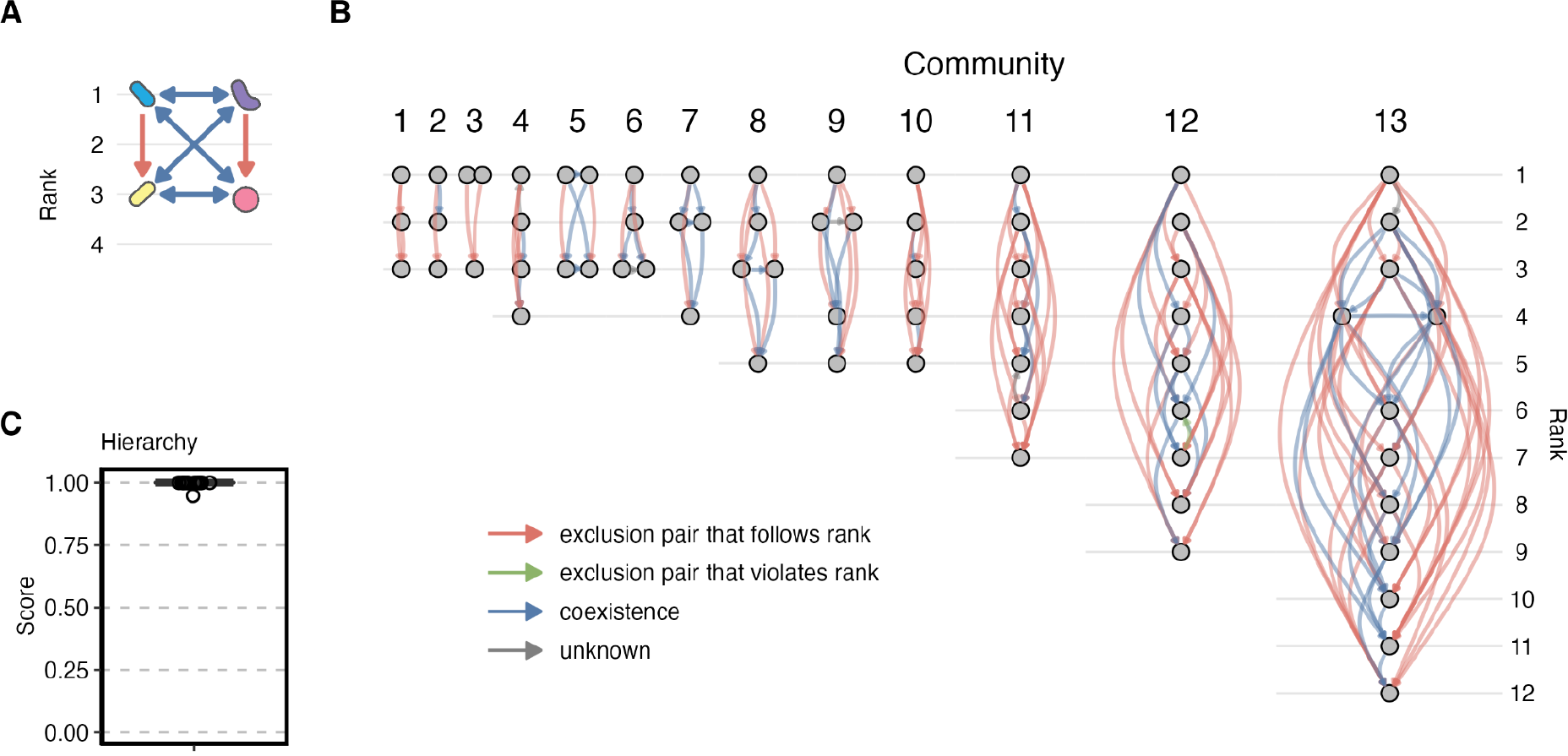
Competitive hierarchy prevails among species pairs in stably coexisting communities. **(A)** Ordering strains by competitive ranking reveals a competitive hierarchy of a representative community (from community C2R6). The direction of the exclusion arrow (red) connects a winner to the loser, whereas the pairwise coexistence links (blue) are mutual. (**B**) The competitive rankings of strains are shown for 13 experimental communities. The gray nodes in the networks denote the microbial strains. (**C**) The hierarchy scores. Each point represents the hierarchy score for an community.

Here, *N_f_* represents the number of exclusion pairs that follow the strains’ competitive rankings (e.g. where the higher-rank species excludes the lower-rank competitor), and *N_v_* represents the number of rank-violating pairs (e.g. where the lower-rank species excludes the higher-rank competitor). Note that this measure only considers exclusion pairs. If exclusion were perfectly hierarchical, we would expect to see *h*=1, as none of the lower-ranked species would exclude a higher-ranked community member. All of the 13 communities in our experiments as well as those in our simulations exhibit a very high degree of hierarchy (*h~1*) (Fig. 4C). Overall, we see no evidence of nontransitivity in our coexistence/exclusion networks.

## DISCUSSION

At the outset of this paper, we set out to empirically address a fundamental question in microbial community ecology, using an empirical system that is uniquely suited for that purpose: Is species coexistence primarily a reductionist pairwise phenomenon, or is it an emergent property of the whole community? To address this question, we reconstituted every possible pairwise co-culture out of the (N=3-12) coexisting members of 13 different enrichment communities, in the same environment where the communities had been assembled, and determined whether the pairs coexisted on their own in the absence of the other members of their communities. We found that they do generally not: pairwise competitive exclusion prevails across these communities. This was true in all of our communities, which exhibit highly similar fractions of coexisting pairs regardless of their diversity. These findings indicate that the community context is essential to explain pairwise coexistence within multispecies communities. Our results also suggest that coexistence cannot be trivially reduced to pairwise coexistence rules, and indicate that pairwise competition experiments where two species are grown in isolation may not be very informative about the ability of those species to coexist, even when done under identical abiotic conditions as those experienced by the community. Our results do not allow us to determine whether the complex nature of multispecies coexistence derives from higher-order interactions, or whether it can be explained by a complex network of pairwise interactions. Additional experiments will have to be done in future work to settle this question.

Several theoretical studies have suggested that nontransitivity stabilizes the coexistence of a large number of competing species when the competitors have differential competitive abilities on multiple limiting resources^4,5^, or in the presence of spatial heterogeneity^9^. Despite the well-established theoretical basis of this idea, empirical studies on the prevalence of interspecific nontransitivity are still sparse. Taking the advantage of the mutual invasion experiments of 186 species pairs from each of the 13 replicate communities, we found strongly hierarchical competition among our species, and found no evidence of nontransitivity, suggesting that the role of nontransitive competition in promoting species coexistence may be negligible in our communities. The discrepancy between the theory and our findings may arise from the underlying ecological interactions among competing species. In our communities, nutrient exploitation on the single limiting nutrient and cross-feeding are the dominant ecological interactions in determining the community structure^29,30^, whereas laboratory microbial experiments have suggested that nontransitivity may emerge through mechanisms such as changes in species’ competitiveness across resources^36^ and the balance among toxin-producing, resistant, and sensitive strains^23^.

Future work should address to what extent the topology of the resource competition and cross-feeding networks may affect the answer to the question raised above. While this is beyond the scope of this paper, the empirical system developed here could be extended to environments containing a larger number of resources, as done elsewhere^31,34,37^. This could be a way to manipulate these networks empirically. In sum, we hope that the methodology developed here will help us determine under what conditions one should expect simple pairwise assembly rules to predict the assembly of microbial communities. Our findings highlight the challenges of predicting species coexistence from a bottom-up perspective and emphasize the importance of developing models that take into account the full community context.

## ACKNOWLEDGEMENTS

We want to thank members of the Sanchez Lab for their helpful discussion. This work was partially funded by Young Investigator Award RGY0077/2016 from the Human Frontier Science Program (to A.S.). C, C.-Y. was supported by the Graduate Student Fellowship for Studying Abroad from the Ministry of Education, Taiwan.

## CONTRIBUTIONS

C.-Y.C., D.B., and A.S. conceived the idea and designed the study. C.-Y.C. carried out all experiments, made the results figures, and made the diagrams. C.-Y.C., D.B., J.C.C.V, S.E., and A.S. discussed the results and drafted the paper. C.-Y.C.. and A.S. wrote the final version of the paper.

## COMPETING INTERESTS

The authors declare no competing interests.

## Notes

### Competing Interest Statement

The authors have declared no competing interest.

### Summary of Updates

Instead of using a hybrid of manual colony counting and sequencing, we have shifted to using an automatic image processing pipeline combined with a machine learning approach to classify bacterial colony morphology. Accordingly, Figures and Supplements are updated.

