## Supplemental Methods, Figures, and Tables for "Emergent coexistence in multispecies microbial communities"

This PDF includes
Methods

- Experimental design
- An automatic pipeline for extracting colony features from scanned images
- Colony morphotype classification using a supervised machine learning algorithm
- Determining the competition outcomes
- Competitive network analysis

Supplementary Figures 1-6
Supplementary Tables 1-3

All the R and Python scripts for analyses and plotting are on GitHub repository
https://github.com/Chang-Yu-Chang/emergent-coexistence

### Methods

#### Experimental design

**Isolation of microbial strains**

Communities used in this study were assembled from various foliage- and soil-derived microbiomes in a minimal medium with M9 salts and 0.2% glucose as the single supplied carbon source. These communities were allowed to equilibrate for 12 transfers (~84 generations) in batch culture. Detailed assembly processes and community properties were described in our previous work [1]. To isolate the microbial strains for pairwise competition experiments, we streaked out the frozen stocks of 13 representative communities on nutrient agar plates (trypticase soy agar; TSA), picked morphologically distinguishable colonies, and re-streaked them on selective agar plates of M9 salts with glucose or citrate as carbon sources. We then sequenced the 16S ribosomal RNA (rRNA) gene of each isolated strain, merge 16S Sanger sequences, and compared the sequencing results to the bacterial 16S rRNA database to determine their taxonomic identities (see below). For storing the stocks of the isolated strains, we dissolved the colonies with phosphate buffer saline (PBS) and mixed them with an equal volume of 80% glycerol, and saved the stocks at -80℃.

**Sanger sequence alignment and taxonomy assignment**

To align and trim the Forward and Reverse reads from Genewiz, we used a function sangerAlignment from the R package “sangeranalyseR” [2]. This algorithm trims both ends of the two sequencing reads with a low quality score using Modified Mott Trimming, then aligns the two trimmed reads as a full-length 16S rDNA sequence. We used the default setting of the sangerAlignment function. Once we obtained the full-length 16S rDNA sequences, we assigned the isolate’s taxonomy using the R tool of the RDP classifier [3,4].

**Matching isolates Sanger sequences to amplicon sequences**

To determine isolates’ relative abundances in the community, we matched the isolates’ full-length 16S rDNA Sanger sequences to the amplicon sequencing data (ESVs) of the communities by performing a pairwise alignment using the function pairwiseAlignment from the R package Biostrings, with alignment type set to ”local" [5]. We aligned the full-length Sanger sequence of an isolate with all possible ESVs from the same community and obtained the alignment scores. We then chose the Sanger-ESV alignment with the highest alignment score. Because the Sanger sequences cover the full-length 16S rDNA whereas ESVs only use the V4 region, it is possible that two Sanger sequences match one ESV. If this is the case, we chose the isolate with the higher alignment score, resulting in 46 unique Sanger-ESV pairs (22 of 68 isolate Sanger sequences were dropped). In the 46 pairwise alignments, the shortest consensus length is 233 base pairs, with 33 full matches, eight one-base-pair mismatches, two two-base-pair mismatches, one three-base-pair mismatch, and two four-base-pair mismatches.

**Calculating the pairwise phylogenetic distance between two competing isolates**

To determine the phylogenetic distance between two competing isolates from one community, we computed the difference in the 16S sequences between two competing strains. This includes two steps. First, we trimmed the 16S sequences based on each base's quality score. Specifically, we used Richard Mott's modified trimming algorithm (PRIVATE), adapted from Biopython.SeqIO [6]. Second, we aligned the trimmed sequences of two isolates and computed the number of base pair mismatches between the two trimmed sequences. We conducted the alignment and base pair comparison using the EMBOSS needle algorithm [7]. The resulting data is a matrix where the rows and columns denote the isolates and the values represent the number of base pair mismatches between the two isolates in comparison.

**Pairwise competition experiments**

We streaked out the frozen stocks of 68 bacterial strains on the nutrient TSA agar plates and grew them for 48 hours at 30℃. The agar plates were then stored at 4℃ for up to two weeks before being used in the competition experiments. On the first day of the experiment, we prepared the strain inoculum by suspending the colonies of each strain in PBS, which were standardized to an optical density (OD) of 0.1 at 620 nm. To initiate the competition at three mixing frequencies, we mixed the inocula of two competing strains by volume to the desired ratios of 5%:95%, 50%:50%, and 95%:5% in 96-well 200 µL nunc plates as a master copy. We then inoculated 4 µL of the inoculum mixtures to 500 µL of M9 glucose liquid medium contained in 2 mL 96-well deep-well plates, which were then sealed with porous film. For each growth cycle, we incubated the cultures at 30℃ without shaking for 48 hours and diluted the mature cultures by a factor of 1/125 into a fresh M9 glucose medium to start the next cycle. We carried out the competition experiments for eight growth cycles and made an experimental duplicate for each of the three mixing frequencies. The duplicate was included in the experimental design as a sanity check and only one replicate was eventually plated on agar plates for determining the final frequencies.

*Experimental batches*. Because of the experimental scale and the limitation on the number of cultures that can be handled at a time, we conducted the experiments in four different batches: B2, C, C2, and D. Batch B2 includes six communities and a total of 27 bacterial isolates. The number in parentheses indicates the number of isolates in each community: C2R6 (4), C2R8 (4), C7R1 (4), C8R4 (3), C10R2 (3), C11R1 (9). Note that in batch B2, the inoculum of isolate 1 from community C11R1 was contaminated, so all cocultures containing this isolate from batch B2 were removed from the analysis. Instead, the competition experiments of these cocultures were replaced by batch C where we only carried out the competition experiments of this isolate with other strains in the same community. Batch C2 contains the largest community C11R2 which has 12 isolates. Batch D has six communities and a total of 34 isolates: C1R2 (4), C1R4 (5), C1R6 (5), C1R7 (7), C4R1 (3), and C11R5 (5).

*The naming convention for cultures and images*. We have a naming convention for colony plating and image processing in order to keep track of the cultures. This naming system appears as the image file names and the sticker labels on the petri dishes (see the Plating and Scanning section below). We designed it in a way that we can easily parse the names using regular expression to extract information such as experimental batch, source community, transfer, mixing ratio, and isolate. For example, an image file of monoculture is named "B2_T1_C2R6_1", which indicates batch B2 at transfer T1 (culture after the first growth cycle), and the strain is community C2R6 isolate 1. For cocultures, an image file named "C2_T8_C11R2_5-95_1_2" means batch C2 at transfer T8 (the last growth cycle), community C11R2, mixing ratio 5:95 between isolates 1 and 2. Note that isolate 1 in this coculture starts at 95% and isolate 2 at 5%. Whereas a coculture named "C2_T8_C11R2_5-95_2_1" means that isolate 1 starts at 5% and isolate 2 at 95%. This naming convention was applied to the file names of the raw images as well as the temporary images.

**Plating and scanning**

*Plating isolate inocula, monocultures, and cocultures.* We plated the inocula and cultures at four time points: the monoculture inocula at the beginning of the competition experiment (monocultures at T0), the monoculture after the first growth cycle, and the last growth cycle (monocultures at T1 and T8), as well as the cocultures after the last growth cycle (coculture at T8). Before the day of plating, we prepared 20 mL of TSA agar medium, which was the same medium used originally to isolate the strains, in the clear sterile 100mm x 15mm polystyrene petri dishes (FisherBrand). Once the agar was properly solid, we wrapped the loaf of agar plates in plastic bags and stored them at 4℃ to keep them moist and fresh before plating. On the day of plating, we serially diluted the culture samples by a factor of $10^{-5}$. Specifically, we transferred 10 $\mu$L of the culture and mix it with 90 $\mu$L of PBS on a clear 96-well nunc plate. We then dispensed 10 $\mu$L of the mixture and mix it with another 90 $\mu$L of fresh PBS, and repeated this step five times until we obtained the $10^{-5}$ dilution. We then dispensed 20 $\mu$L of the $10^{-5}$ diluted sample onto a TSA agar plate, added 5-7 sterile glass beads, and shook the plate until the liquid culture was evenly smeared on the surface of the agar. We removed the beads and placed the inoculated petri dishes at 30℃ for three days.

*Scanning setting*. After three days of incubation when the bacterial colonies developed to a noticeable size, we scanned the petri dishes using a scanner model with a transparency unit (EPSON Perfection V850 scanner). We connected this scanner to a modern Mac and configure the scanner parameters in the compatible software ImageCapture (Scan Mode: Transparency-Positive, Kind: Color Slide, Colors: Billions, Resolution: 600 dpi, Format: TIFF). To ensure a consistent image dimension, we used a fixed custom scan size by setting a square of edge length 3.7 inches as the scan region. To avoid the droplets and reflections that may obscure the scan, we removed the petri dishes' lids before placing them on the scanner. Once scanned, image files were named to match the labels on the petri dishes. It is possible that incubation time may affect the colony morphology as the longer the incubation time the larger the colonies develop. To minimize this factor that may affect the morphology identification, we always scanned the coculture petri dishes from the same experimental batch within a day.

### An automatic pipeline for extracting colony features from scanned images

Once we obtained the scanned images of the plates, we employed an automatic pipeline that detects the colonies and returns their features such as shape, size, and textual pattern (Fig. S1). Briefly, this pipeline includes three major parts: color management, image segmentation, and feature extraction. Most steps were executed under the R environment using an R package "EBImage" [8] except for the background subtraction where we used the Python package "scikit-image" [9]. These steps are wrapped in independent scripts that allow consecutive execution of multiple images in a command-line environment.

**(I) Color management**

*(1) Grayscale of RGB channels.* The original images are red-green-blue (RGB) raster images stored in a TIFF format with dimensions width $\times$ height $\times$ channel = 2220 $\times$ 2220 $\times$ 3. In this step, the three color channels were split and displayed in grayscale. Specifically, we read the TIFF image in R, set the color mode of the RGB image to grayscale, split the color channels, and stored the three greyscaled images in separate folders in the TIFF format. We executed these steps using readImage, colorMode, and writeImage functions from EBImage.

*(2) Background subtraction*. To correct for unevenly illuminated background introduced by the agar medium, we implemented background subtraction using a "rolling ball" algorithm. This algorithm calculates the background value at the desired pixel by taking into account the intensities of its surrounding pixels. The size of the local region depends on a key parameter "radius". When the radius is small, only a small number of neighboring pixels are considered and therefore more details of the local background are conserved; whereas when the radius is large, the details are smoothed out. We executed background subtraction using the rolling_ball function from the Python package scikit-image with the radius set at 80.

**(II) Image segmentation**

Segmentation is the process of partitioning a digital image into multiple image segments, or objects. In our case, the target objects are bacterial colonies. Here we carried out segmentation in five steps: thresholding, brushing, first object filter, watershed, and second object filter. To maximize the color contrast between the objects and background for better image segmentation and reduce computational loads, we chose the green channel for segmentation as by eyes the colonies stand out more clearly from the background in this channel compared to the other two channels. Therefore, only the green-channel image was segmented. After segmentation, we used the segmented green-channel image as a roadmap to extract the textual features in all three channels.

*(3) Thresholding*. The first step of segmentation is thresholding, which sets pixels whose values are above a threshold to a foreground value and all the remaining pixels to a background value, resulting in a binary image. To further account for spatial variation in illumination, instead of a global threshold, we adopted an adaptive thresholding approach that compares the image with its filtered version from a moving

rectangular window. We implemented this step using the thresh function from EBImage with a moving window of width=150 and height=150, and the offset value set at 0.1. The returned binary image was then passed to the brushing step.

*(4) Brushing*. Sometimes the particles or uneven illumination in the background may result in a large number of tiny objects that do not resemble the target objects (in our case the colonies). These off-target objects increase unnecessary computation time. To remove such background noise, reduce the computation loads, and smooth the object contour, we implemented a brushing procedure that includes three steps. First, we generated a disc-shaped “paintbrush” with a diameter=11. Second, we put this paintbrush over every foreground pixel (pixels of the objects) in the binary image, setting the focal pixel to the background value if any of the pixels covered by the paintbrush is from the background. In other words, the objects are said to be “eroded” after this step, and any objects smaller than the paintbrush would be set to the background. Third, we put the paintbrush over every background pixel (pixels of the background) and set the pixel to the foreground if any of the pixels covered by the paintbrush is from the foreground. This step is called “dilation” as the contour of the large objects is smoothed. We generated a disc-shaped paintbrush using the makeBrush function with the size set to 11 and the shape set to “disc”, and implemented the erosion and dilation steps using the opening function.

*(5) First object filter*. To further remove the non-colony objects before the computationally demanding watershed object detection, we applied an arbitrary filter according to the object size and its position in the image. First, the object size has to be higher than 300 and lower than 20000 pixels. This range was determined after a few trials such that it covers almost every object with comparable size as the colonies while removing objects that are obviously too large or too small for a colony. Second, the objects have to be located within the agar medium of the petri dishes. One major source of false-positive objects comes from the edges of the petri dishes where a colony cannot possibly occur (Fig. S2A). To address this issue, we removed an object if any of its contour pixels fall out of the range [100, 2120] in both the x and y coordinates, meaning that this object is too close to the image edges (within 100 pixels) that could not possibly be a colony.

*(6) Watershed object detection*. When two colonies grow too close to each other, they would eventually fuse and may be falsely detected as one object instead of two. We addressed this problem using a watershed algorithm to segment the fused colonies into two or multiple objects. First, the input binary image is transformed into a distance map which is a matrix containing for each pixel the distance to its nearest background pixel. Second, the watershed algorithm inverts the distance map and uses water to fill the resulting valleys (the center of a tentative object) until the filled water from another object or background is met. We calculated the distance map using the distmap function and implemented the watershed algorithm using the watershed function from the EBImage package. After the watershed, the resulting image has each object labeled, meaning that each labeled object is a set of pixels with the same unique integer value.

*(7) Second object filter*. After the watershed, sometimes false-negative objects might appear as multiple colonies form a clump and the watershed algorithm fails to distinguish them apart (Fig. S2B). We applied a second filter to further remove such colony clumps according to object size and shape. First, the object size has to be larger than 300 pixels, which is the same criterion for the minimal colony size as described

in the first object filter. Second, we filtered for the objects with circularity $C>0.7$. Circularity is defined as $C=4\pi AL^2$, where A is the object area and L is the perimeter. The circularity C ranges in between [0,1]. Any shape that is extremely close to a line has C=0, whereas a perfect circle has C=1. Third, the ratio between the standard deviation of the radius and the mean radius has to be lower than 0.2. This filter removes those objects that are roughly round but has a very rugged contour (for instance a clump with many small colonies) which the circularity filter may fail to detect.

**(III) Feature extraction**

*(8) Computing object features.* Computing the object features requires two input images: a roadmap image and a reference image. The roadmap image contains labeled objects, which are pixel sets with the same unique integer value. Such roadmap images are the resulting images from the image segmentation step described above. The reference image contains the intensity values of the reference objects. In our case, the reference images are the three color-channel images resulting from color management. For each grayscale image from the three color channels, we computed morphological and texture features such as radius and means intensity using the computeFeatures function from EBImage, as well as the transect features using a custom function calculate_transection (see the section below), resulting in a table of object features. In this table, rows represent the detected objects and columns denote the object features. For one image file, we stored features obtained from the three color channels in a CSV format. For each object, we obtained a total of 40 features, where 6 are shape-related, 5 are moment-related, 23 are intensity-related from the green channel, 3 are intensity features related to the red channel, and 3 are related to the blue channel (Table S1).

*Transect features*. Some colonies develop distinct patterns of radial gradient that serve as a reliable cue for distinguishing two morphotypes. Radial gradients are radial changes in pixel intensity from the center to the edge of the colony. For example, the colonies may display noticeable outer rings, concentric circles, or gradual changes (Fig. S2D-H). To quantify this observation as the object features, we drew a transect for an object and computed the intensity values along the transect. Specifically, in the watershed image, we calculated all radii of an object by computing the distance between the center and each contour pixel. We then searched for the contour point with the median radius, drew a transect between this contour point and the object center, and locate all pixels covered by this transect (transect pixels). We extracted the intensity values of the transect pixels from the reference image (only the green channel image). For an object transect, we calculated the summary statistic of pixel intensity such as mean, standard deviation, and quantiles (Table S1). To capture the observed radial gradient as the colony features, we also divided the transect into segments and extracted intensity from the pixels at the breakpoints. Specifically, we re-scaled the transect length into a unit of 1 and located the pixels that are 0.05, 0.1, 0.5, 0.9, and 0.95 units away from the object center.

*Intensity feature anomalies*. Sometimes a colony may be located adjacent to the plate edges, where the colony is shaded and therefore has a high variation in pixel intensity (Fig. S2C). To remove these anomalies, we applied an outlier filter where seven intensity variation features were considered: "b.sd", "b.mad", "b.mean", "b.q05", "b.q005", "b.tran.sd", and "b.tran.mad" (Table S1). Specifically in each segmented image, we calculated the lower quartile ($Q_1$), the upper quartile ($Q_3$), and the interquartile range (IQR) of a targeted feature. An object is deemed as an outlier if its feature value falls outside the range of [$Q_1$ - 2 × IQR, $Q_3$ + 2 × IQR] and is then removed. We repeated the same rule for each of the seven intensity features.

In summary, we have 68 monoculture images and 558 coculture images. Among 186 species pairs, we removed six pairs where the coculture images do not have any colony, resulting in 180 unique species pairs entering the classification step below.

#### Colony morphotype classification using a supervised machine learning algorithm

Our goal is to classify the coculture colonies based on the monoculture colony morphology. To achieve this goal, we adopted a machine-learning approach. In particular, we used the Random Forest algorithm, a widely used supervised machine learning method for classification and regression problems. In short, for each coculture image, we trained the model using the labeled colony feature data from monocultures. The training procedure involved cross-validation in tuning the hyperparameter according to the model accuracy. Once the hyperparameter was determined, we then fit the model using the monoculture dataset and used this model to predict the colony morphotype on the coculture image. We repeated this procedure for all 180╳3=540 coculture images. The monoculture and coculture images, Random Forest result, and object feature scatterplots for each of the 540 cocultures are shown in four Supplementary Material PDFs (random_forest-B2.pdf, random_forest-C.pdf, random_forest-C2.pdf, and random_forest-D.pdf). One example is shown in Fig. S3. We performed the Random Forest model training and cross-validation using R packages “randomForest” [10] and “caret” [11].

*Training and validation datasets*. For each coculture, we trained a Random Forest model using the object features (colony features) from each of the two monocultures. First, we read the colony features of both monocultures obtained from image processing, labeled their sources (morphotype A or B), and performed hyperparameter tuning with cross-validation (see the next section). Once we settled on the value of the hyperparameter, we fit the model with this hyperparameter using the monoculture colony morphology features. We then used this model to predict the coculture colony morphotype. Regarding the source of images, we used T8 images (cultures after the last growth cycle) for both the monoculture and the coculture images. For some monocultures, we used the T0 (inoculum) or T1 (cultures after the first growth cycle) images instead, either because the strain did not grow on the minimal medium therefore no colony was present at T8 (for instance, community C11R1 isolates 2, 8, 9), or because there is cross-contamination where the T8 colony morphology looks noticeably different from that at T0. The list of isolates and their corresponding monoculture image and colony count are shown in Table S2.

*Tuning model hyperparameter*. One key hyperparameter in Random Forest is MTRY, which specifies the number of features that the algorithm can select from at a tree node split. If MTRY is too small, during the tree-growing process, unimportant features may be selected multiple times which reduces the model accuracy; whereas if MTRY is too large, in the extreme scenario, equal to the total number of all possible features, the randomness would vanish because the most important feature is selected every single time the node splits. Taking this into consideration, we performed repeated cross-validation across a range of eight MTRY values (2, 7, 12, 17, 22, 27, 32, and 37) and chose the MTRY value with which the model accuracy peaks. Specifically, we performed three repeats of 10-fold cross-validation, meaning 30 training/validation sets for each MTRY value. For each training/validation set, the model accuracy was calculated as the proportion of the classified objects that are correct in the validation set. We then computed the mean accuracy across the 30 training/validation sets for each MTRY value and chose the MTRY with the highest mean accuracy. Another key hyperparameter NTREE which specifies the total number of decision trees is always set to 500.

*Model accuracy and prediction*. We performed independent hyperparameter tuning and modeling training for each of all 540 coculture images. In some cocultures, the two morphotypes may resemble each other so the model failed to distinguish them apart. Taking this into account, we set an arbitrary filter at accuracy = 0.9 as the criterion for quality control of the model training. Following this criterion, we discarded 9 of 180 unique species pairs, resulting in 171 pairs for the later analysis (Fig. S4). With the trained model, we were able to use the colony features to predict the colony morphotype occurring in the coculture images. The Random Forest model accuracy and object prediction for each of the 540 coculture images are shown in the four Supplementary Material PDFs.

*Comparing human and machine results.* To double-check the efficacy of our machine-learning approach, we compared the machine results with human results. In the human results, the first author manually counted the colonies from the images by eye. The comparison between human and machine results is shown for the total colony counts and the morphotype frequency (Fig. S5). In general, the machine-detected total colony counts correspond to the human results (linear regression R-squared=0.88), suggesting that the image segmentation successfully detects colony objects. In the case of morphotype frequency, we observed a statistically significant correlation between the human and machine results (linear regression; adjusted R-squared = 0.85).

### Determining the competition outcome

To determine whether two competing strains coexist stably in a pairwise competition experiment, we considered an isolate's fitness function, which determines whether the focal isolate can increase when rare and decrease when abundant. The fitness functions can be measured by comparing the isolate's three initial frequencies between the beginning and the end of the competition. To obtain this data, there are two issues to be addressed. First, the initial frequencies were manipulated using OD (5:95, 50:50, and 95:5) whereas the final frequencies of coculture can only be measured using CFU instead of OD. Unfortunately from hindsight, we did not plate the T0 coculture inoculum so the CFU frequency data at T0 is unavailable. To address this discrepancy in data types, we converted the OD to CFU for the initial inocula using the monoculture dataset. Second, each time we plate the colonies from diluted culture, we essentially "sampled" dozens to hundreds of cells only once from a pool of billions of cells. Taking this into consideration, we performed bootstrapping on the CFU frequencies at both T0 and T8, where we resampled each colony from the Poisson distribution parameterized by the observed colony occurrence probability.

*OD to CFU conversion at T0.* We calculated the OD to CFU conversion using the monoculture dataset that contains both OD and CFU for an isolate (Table S2). From these plates, we segmented the images using our automatic image processing pipeline to obtain the colony count per image. Typically, we used monoculture data at T8. Because some isolates did not grow in isolation on the minimal medium, their monocultures at T8 do not have any colony. If this occurred, we used the T0 or T1 plates for these isolates. The OD data are either from the set OD=0.1 for T0 or from the microplate reader (ThemoFisher Multiskan FC) measurements at the end of each growth cycle for T1 and T8. From this dataset, we calculated the conversion coefficient $\varepsilon = CFU / (OD_{620}\ \psi\ v)$, where CFU is the number of observed colonies, $OD_{620}$ is the measured OD at 620 nm, $\psi$ is the dilution factor, and v is the volume taken from the diluted culture for platting. We had $\psi=10^{-5}$ and v=20 $\mu$L across all plating practices. The conversion coefficient $\varepsilon$ is strain-specific and invariant to any given dilution factor.

Once we obtained $\varepsilon$ for each isolate, we calculated the hypothetical strain CFU in a coculture at T0. For instance, if in a coculture, isolate A and isolate B are mixed at 5% and 95%, respectively, the hypothetical CFU for isolate 1 is then $CFU_{A,T0} = OD_{620}\ p_A\psi_h\ v\ \varepsilon_A$, where $p_A$=0.05 is the mixing ratio of isolate A. To avoid a low $CFU_A$ that may result in zero counts when used to parameterize a Poisson distribution, we set the hypothetical dilution factor $\psi_h = 10^{-4}$ instead of $10^{-5}$. The T0 inocula were experimentally set at an optical density of $OD_{620}$ = 0.1 and the plating volume v=20 $\mu$L. After plugging these values in the calculation, we retrieved the hypothetical $CFU_{A,T0}$ . Similarly, we repeated the conversion for isolate B with $p_B$=0.95 and $\varepsilon_A$ to obtain $CFU_{B,T0}$.

*Bootstrapping CFU frequencies.* To determine whether the initial frequency (e.g., 5%) of isolate A in coculture significantly increases or declines after competing with isolate B, we bootstrapped both T0 and T8 CFU frequencies 1000 times. To initial bootstrapping at T0, we parameterized two independent Poisson distributions PoisA and PoisB with $CFU_{A,T0}$ and $CFU_{B,T0}$, respectively. $CFU_{A,T0}$ and $CFU_{B,T0}$ are the hypothetical CFUs calculated from the previous section. In one bootstrapped sample, we drew one value $n_{A,T0}$ from PoisA and one value $n_{B,T0}$ from PoisB. The CFU frequency of isolate A at T0 in one bootstrap sample is therefore: $F_{A,T0} = n_{A,T0} / (n_{A,\ T0} + n_{B,T0})$. Likewise, we performed bootstrapping at T8 by parameterizing two independent Poisson distributions with the random-forest-predicted CFU of each morphotype, resulting in the CFU frequency of isolate A at T8: $F_{A,T8} = n_{A,T8} / (n_{A,T8} + n_{B,T8})$. The two sets (T0 and T8) containing 1000 bootstrapped samples were then compared in pairs to determine the statistical significance of frequency changes.

*Computing statistical significance*. The frequency changes were considered statistically significant if more than 950 of the 1000 bootstrap samples (p=0.05) have the same sign of change. For example, if one strain starting at 5% in a coculture consistently increases 965 times and decreases 35 times among 1000 bootstraps, it means the "increase" is statistically significant with a p-value=35/1000 = 0.035. When one starts at 5% in coculture, and neither its increase nor its decreases at T8 exceeded 950 times, it suggests that the frequency change is insignificant. We repeated this calculation for the three mixing frequencies for each competing pair of isolates. We then obtained a fitness function specific to each pair, dictating whether a focal isolate would increase, decrease, or stay at a similar frequency when it starts at rarity, medium, or abundance when competing with another strain.

*Fitness function and competition outcomes*. With the pair-specific fitness function, we followed a mutual invasion criterion to determine whether the pairwise competition outcomes are stable coexistence. Generally, two species can stably coexist when there is negative frequency dependence in the fitness function. To systematically characterize all possibilities of frequency dependence as well as competitive exclusion, we described the relationship between the fitness functions and the competition outcomes. First, each of the three initial frequency treatments (5%, 50%, or 95%) has three possible signs of change (increase, decrease, or non-significance), giving rise to 27 possible fitness functions (Table S3). Among these 27 fitness functions, there are five possible competition outcomes: exclusion, frequency-dependent coexistence, and coexistence at a low abundance (lack evidence of frequency-dependent coexistence), neutrality, and unknown. We explain the criterion for each competition outcome below.

*Competitive exclusion*. A pair was considered as competitive exclusion when the frequency of one species always decreased regardless of its initial frequencies, and we determined the competitive dominance by connecting the competitively superior to the inferior species. Mutual exclusion occurred whenever a strain starting from abundant always excluded its rare competitor and vice versa. Among the 27 fitness functions, two represent competitive exclusion (1 and 14), and another two indicate mutual exclusion (10 and 13).

*Frequency-dependent coexistence*. Frequency-dependent coexistence occurs whenever one species always increases in frequency from rare and vice versa, leading to coexistence at an equilibrium. This can occur when there is a globally stable equilibrium (2, 5, and 8) or when the frequency starts closer to equilibrium otherwise leads to exclusion or neutrality (4, 6, 11, and 20).

*Possible coexistence at a rare-abundant ratio*. There are cases when a focal strain starts from the rarity and stays at a similar figure from abundance. This may arise because the actual stable equilibrium is close to a low abundance (e.g., at 5% or at 95%). Despite that there is no direct evidence of frequency dependence, we still considered this type of fitness function as coexistence because the rarer species did not go extinct. Such examples are fitness functions 3 and 23 (Table S3).

*Neutrality*. In these cases, we detected that for one isolate pair, more than two of its cocultures show non-significance in frequency changes between T0 and T8. This suggests that the coculture still have both strains persisting at the time when we ended the experiment. Despite it may seem to suggest pairwise coexistence, we do not have the statistical power to rule out the possibility that one species of this pair may eventually go extinct if we had run the experiment longer. We considered neutrality as coexistence in a broad sense despite that it may not be stable over the long term.

*Unknown competition outcomes*. Asides from the competition outcomes described above, there are seven fitness functions (7, 12, 15, 16, 17, 19, and 22) that we cannot assign to any competition outcomes described above. We marked these fitness functions if any of them is realized.

Finally, we matched the experimental fitness functions of 171 species pairs to the scenario described above to determine their competition outcomes (Table S3). For simplicity in the network analysis, we also classified these competition outcomes into three broad groups: exclusion, coexistence, and unknown. The exclusion group includes the clear case of competitive exclusion (1, 14) and mutual exclusion (10, 13). The coexistence group includes the clear case of stable coexistence (2, 5, 8), frequency-dependent coexistence (4, 6, 11, 20), possible coexistence at a low abundant (3, 23), and neutral coexistence (9, 18, 21, 24, 25, 26, 27).

#### Competitive network analysis

To compute and illustrate the competitive hierarchy of communities, we built a competitive network for each community. In a competitive network, each node represents an isolate and the edges are the competition outcomes. For the sake of simplicity, we only considered three coarse-grained outcomes: exclusion, coexistence, and the unknown as possible edges.

*Building a competitive network for each community*. Building a network requires two datasets, one for the node characteristics and another for the edge characteristics. The node dataset includes the isolates' identity, whereas the edge dataset contains the pair of isolates, their competition outcomes, and the direction of the competitive hierarchy.

*Computing the isolate's competitive score*. To rank an isolate by its competitive ability, we computed its competitive score accounting for how many times it wins ($g_w$), loses ($g_l$), or has a tie, and the number of games it has played ($n_g$). The competitive score is then ($g_w$- $g_l$)/$n_g$. Because for some pairs we do not have the competition outcomes due to the absence of colonies or low model accuracy, we removed those pairs from the game. Therefore, within a community, an isolate that has all games included would have $n_g$=n, where n is the number of isolates in the community, whereas an isolate that misses one competition result has $n_g$=n-1. The isolates are then ranked according to their competitive score, where the top-ranking (rank 1) isolate has the highest competitive score.

### Supplementary Figures

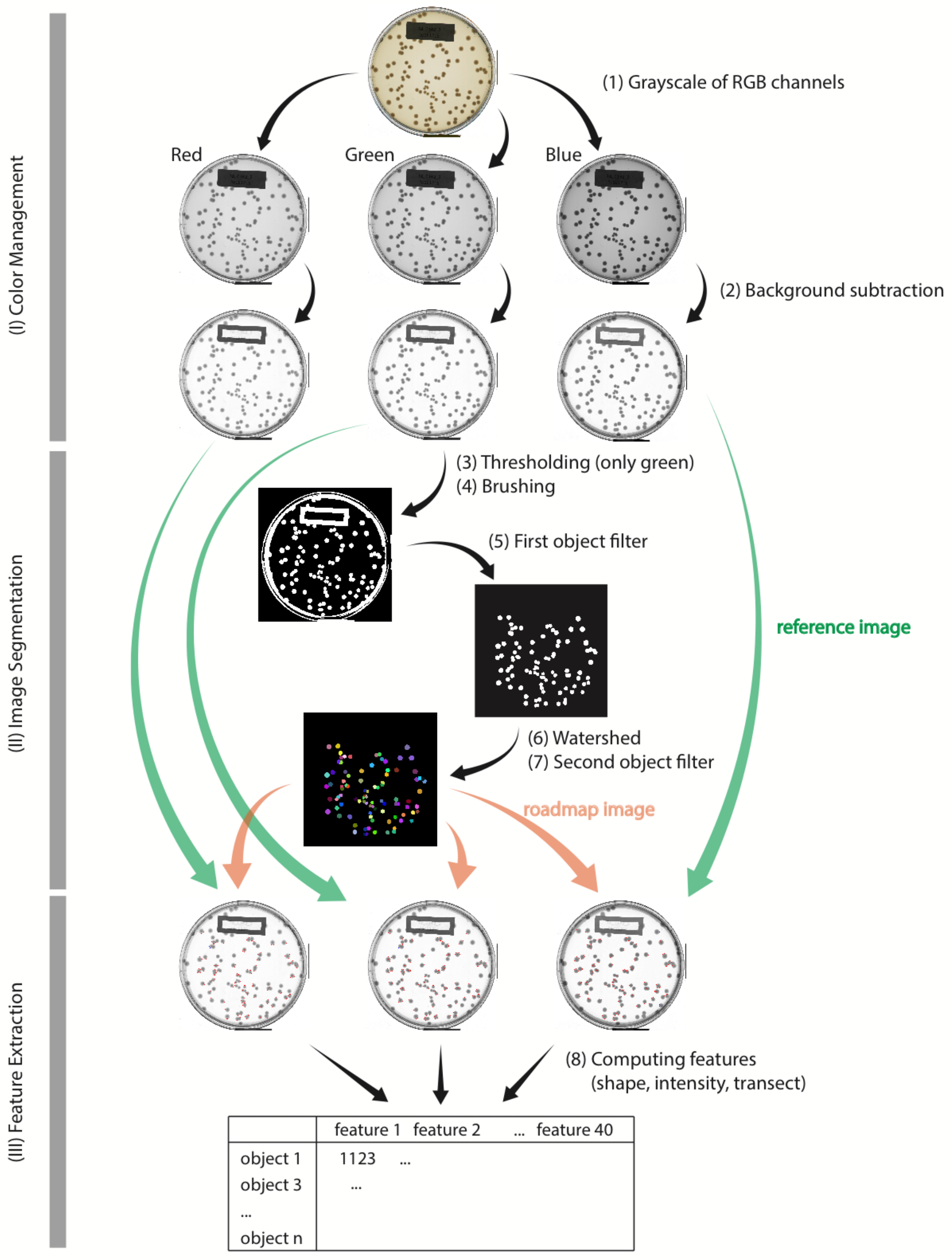


**Supplementary Figure 1. An automatic pipeline for image processing and computing colony features.** This figure shows an example of one image being processed in the pipeline. The main steps (1)-(8) are described in the Methods section.

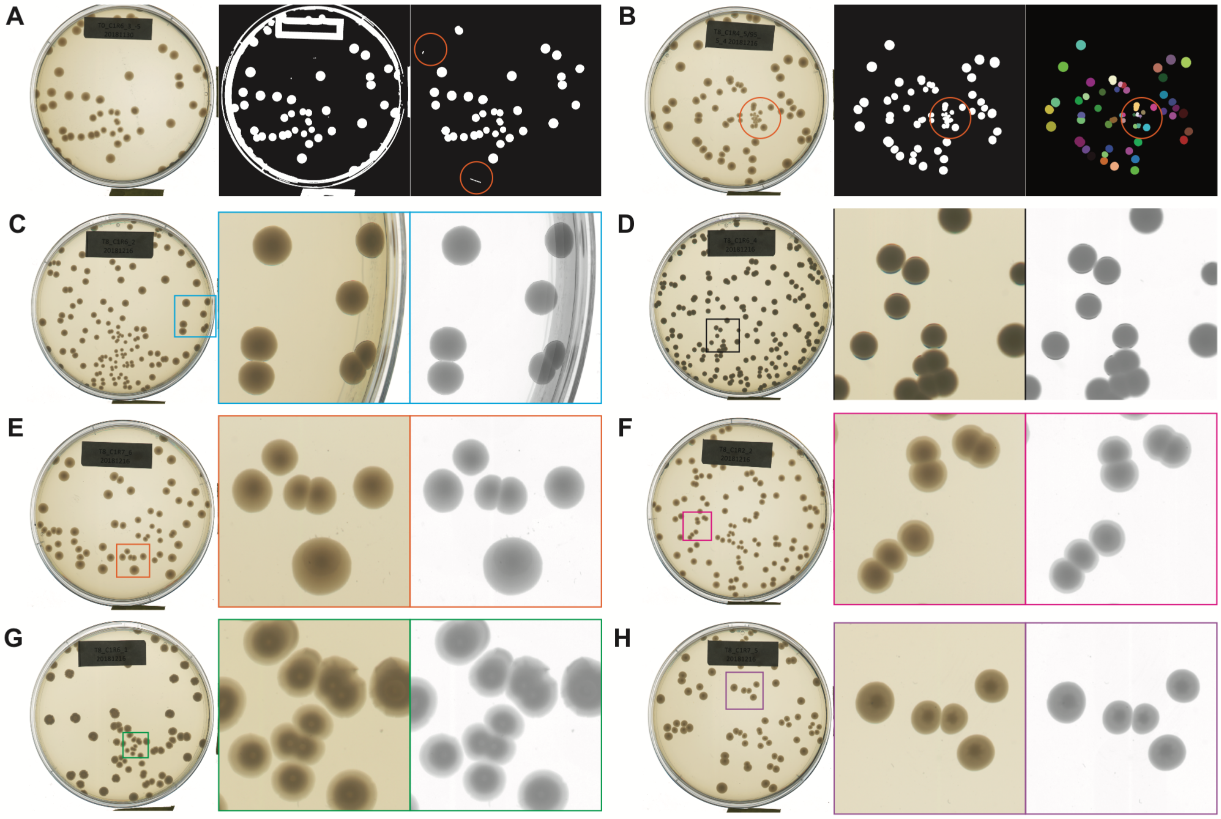


**Supplementary Figure 2. Examples of colony morphologies.** (A) Sometimes after segmentation, an object from the plate edge may be mistakenly identified as an object. These false-positive objects are removed with a second filter with a more strict criterion on object shape after the watershed. The binary images are the thresholded images before and after applying an object filter. (B) Colony clumps occur when the colony density is very high. A clump can have very different shape and intensity properties than the rest of the conspecific colonies, which may mislead model training during the classification. We remove such cases with a more stringent criterion on object shape after the watershed as well. The images shown are before and after the watershed. (C) In some cases, a colony may appear close to the edge of the plate. When it happens, this colony is shaded which largely changes the pixel distribution. We remove these colonies by applying a filter that takes into account the outliers in seven metrics of variation in pixel intensities. A couple of colony morphology examples are shown in (D-H). The most common cases happen when the colony either has (D) homogeneous pixel intensity or (E) a gradual change in the radial intensity pattern. Besides these two cases, some colonies develop distinct patterns, for instance, (F) a light outer ring, (G) concentric circles, or (H) a dark star-shaped center. These features provide key evidence for distinguishing two morphotypes. To capture these features, we compute the pixel intensity change over the transect.

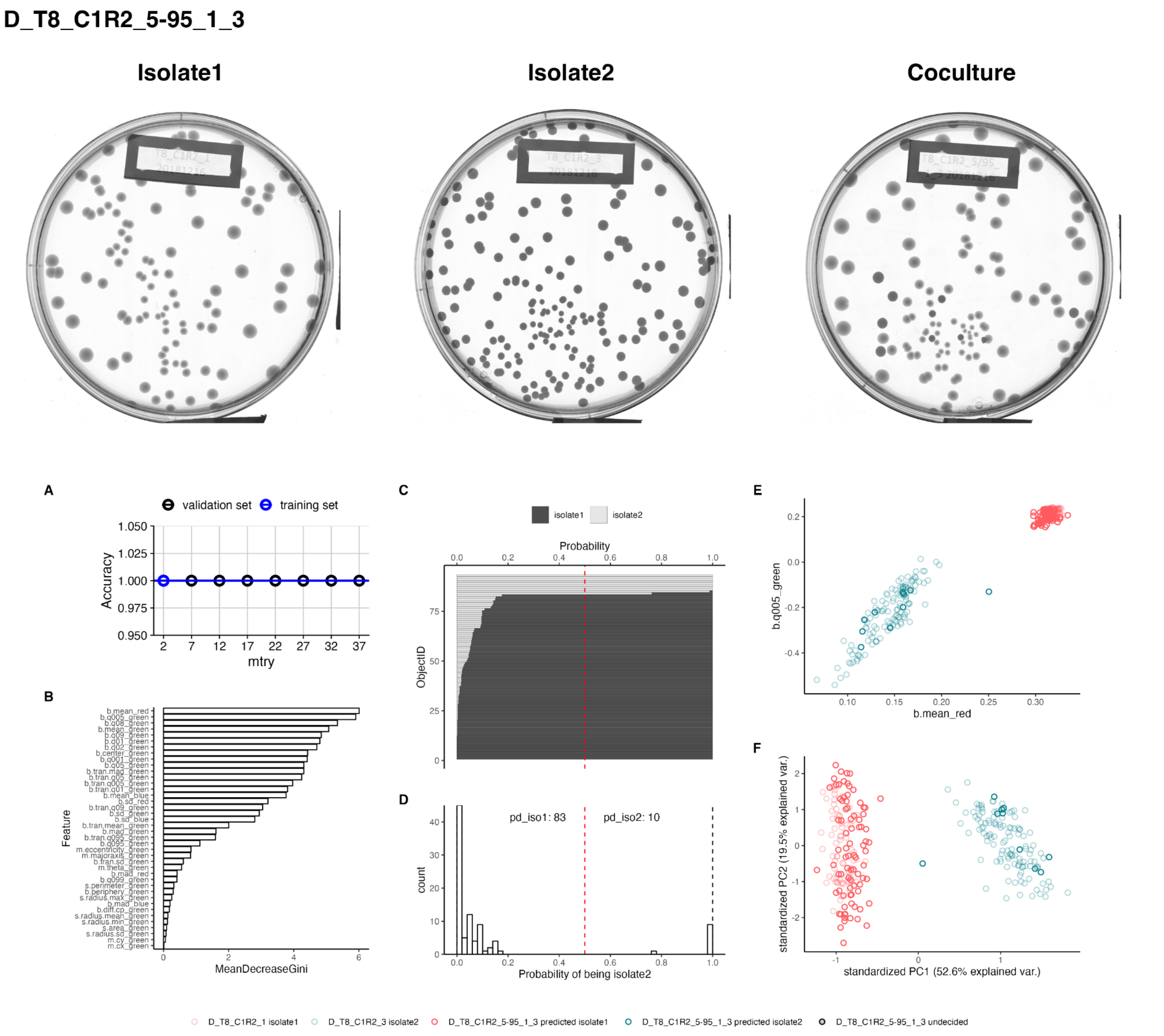


**Supplementary Figure 3. An example of coculture classification using Random Forest.** This figure as an example illustrates one of the 540 coculture classification results included in four Supplementary Material PDFs (random_forest-B2.pdf, random_forest-C.pdf, random_forest-C2.pdf, and random_forest-D.pdf). The top row shows the green-channel, background-subtracted images of two monocultures and the coculture, where Isolate 1 in this case represents C1R2 strain 1, and Isolate 2 represents C1R2 strain 3, and the coculture is when Isolate 1 is mixed at 95% and Isolate 2 at 5%. The lower panels show the Random Forest outputs: (A) the accuracy reported from testing the hyperparameter MTRY at eight values during repeated cross-validation.  (B) the 40 colony features are ordered according to their importance. (C)-(D) For each object segmented in the coculture image, the predicted probability from Random Forest. (E) scatterplot of objects using the top two important features. Light red points denote colonies from Isolate 1 (C1R2 strain 1) monoculture image, whereas light green dots denote Isolate 2 (C1R2 strain 3) monoculture. The dark red and green points represent the predicted morphotype of colonies from the coculture, respectively. (F) a similar scatterplot using instead the top two Principle Components.

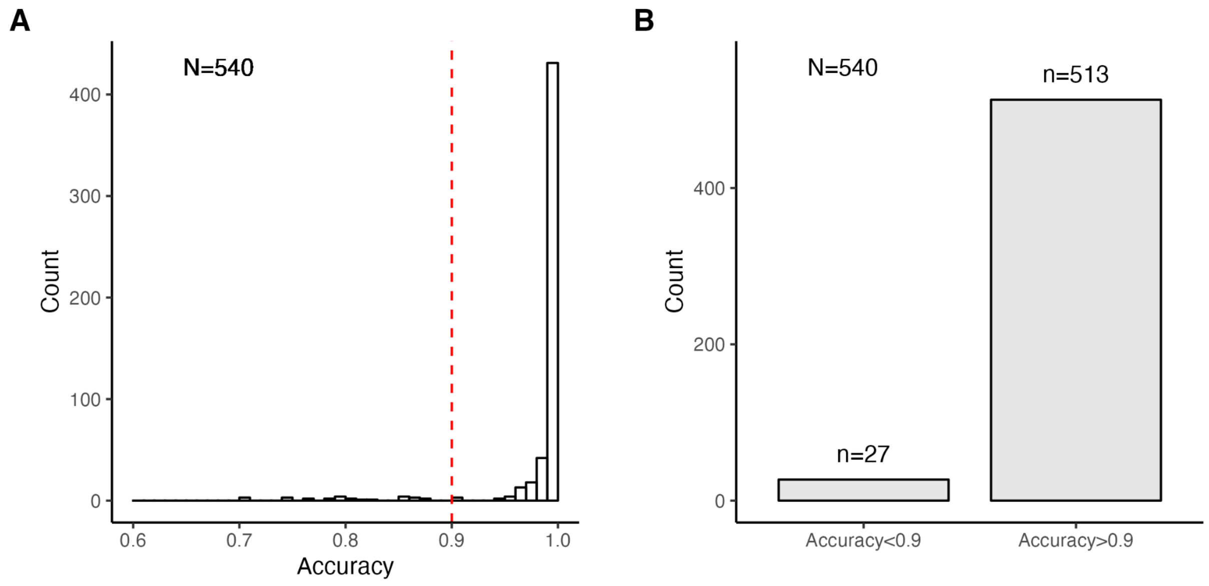


**Supplementary Figure 4. The Random Forest model accuracy for coculture images**. The model accuracy is the fraction where the trained model performs correctly in predicting the object in the validation dataset. Each accuracy value for a coculture reported here was obtained by calculating the average across 30 validation sets. The total number is 180 species pairs and each has three cocultures. (A) the accuracy of all 180✕6=540 coculture images. (B) 9✕3=27 coculture pairs with a model accuracy lower than 0.9 are removed from the analysis.

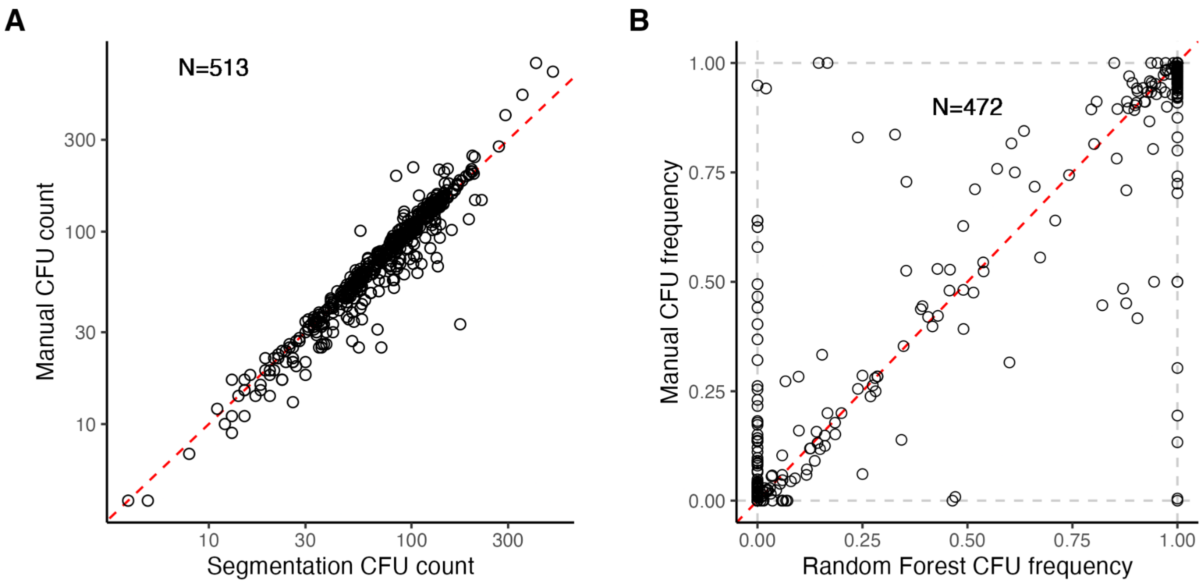


**Supplementary Figure 5**. **CFU results between machine and human results**. (A) The comparison between the CFU count by image segmentation and human counts ($R^2 = 0.8794$). The total number of incidents comes from 171×3=513 after removing 15 species pairs that have no colony in the coculture image or low model accuracy. (B) Random Forest predicted CFU frequency versus the manual results ($R^2 = 0.8499$). The total number of incidents is after removing 41 cocultures where the colony morphology is challenging to human eyes. The dashed red line represents y=x.

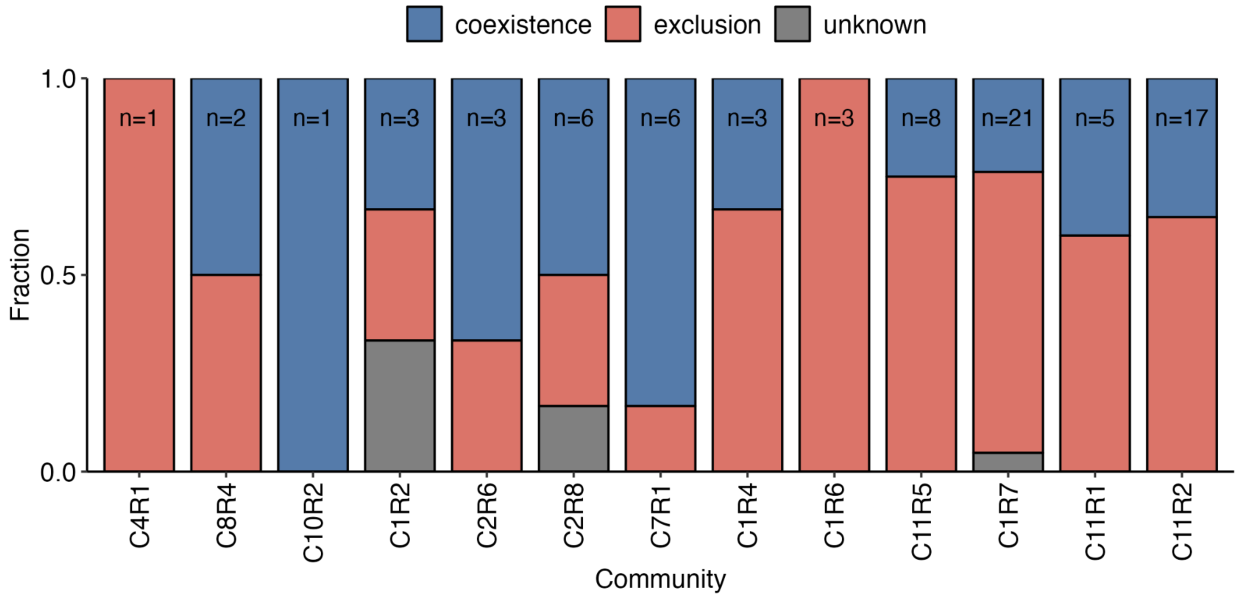


**Supplementary Figure 6. Pairwise competition between highly abundant strains across the 13 communities.** An isolated strain is considered highly abundant in its original community if its matched ESV abundance is above 5%. We consider a pair inapplicable if any of its two competing strains poorly match any ESV or if the strain's ESV is rare. We removed N=92 pairs that meet this criterion, resulting in N=79 pairs where both competing strains are highly abundant. Exclusion remains the most abundant outcome (47/79=59.5%), while coexistence is less common (29/79=36.7%), and three cases being unknown (3/79=3.8%).

### Supplementary Table

**Supplementary Table 1. Object features.** 40 colony object features are detected from the image processing pipeline and considered in the random forest classification.

| | Feature | Feature type | Description |
|---|---|---|---|
| 1 | s.area_green | shape | area size (in pixels) |
| 2 | s.perimeter_green | shape | perimeter (in pixels) |
| 3 | s.radius.mean_green | shape | mean radius (in pixels) |
| 4 | s.radius.sd_green | shape | standard deviation of the mean radius (in pixels) |
| 5 | s.radius.min_green | shape | min radius (in pixels) |
| 6 | s.radius.max_green | shape | max radius (in pixels) |
| 7 | m.cx_green | moment | center of mass x (in pixels) |
| 8 | m.cy_green | moment | center of mass y (in pixels) |
| 9 | m.majoraxis_green | moment | elliptical fit major axis (in pixels) |
| 10 | m.eccentricity_green | moment | elliptical eccentricity defined by sqrt(1-minoraxis^2/majoraxis^2). Circle eccentricity is 0 and straight line eccentricity is 1. |
| 11 | m.theta_green | moment | object angle (in radians) |
| 12 | b.mean_green | intensity | green channel mean intensity |
| 13 | b.sd_green | intensity | green channel standard deviation intensity |
| 14 | b.mad_green | intensity | green channel mad intensity |
| 15 | b.q001_green | intensity | green channel 1st quantile intensity |
| 16 | b.q005_green | intensity | green channel 5th quantile intensity |
| 17 | b.q01_green | intensity | green channel 10th quantile intensity |
| 18 | b.q02_green | intensity | green channel 20th quantile intensity |
| 19 | b.q05_green | intensity | green channel 50th quantile intensity |
| 20 | b.q08_green | intensity | green channel 80th quantile intensity |
| 21 | b.q09_green | intensity | green channel 90th quantile intensity |
| 22 | b.q095_green | intensity | green channel 95th quantile intensity |
| 23 | b.q099_green | intensity | green channel 99th quantile intensity |
| 24 | b.tran.mean_green | transect | green channel transect mean intensity |
| 25 | b.tran.sd_green | transect | green channel transect standard deviation intensity |
| 26 | b.tran.mad_green | transect | green channel transect mad intensity |
| 27 | b.center_green | intensity | green channel the inmost transect pixel intensity |
| 28 | b.periphery_green | intensity | green channel the outmost transect pixel intensity |
| 29 | b.diff.cp_green | intensity | green channel difference between b.center and b.periphery |
| 30 | b.tran.q005_green | transect | green channel pixel intensity at the 5% on the scaled transect |
| 31 | b.tran.q01_green | transect | green channel pixel intensity at the 10% on the scaled transect |
| 32 | b.tran.q05_green | transect | green channel pixel intensity at the 50% on the scaled transect |
| 33 | b.tran.q09_green | transect | green channel pixel intensity at the 90% on the scaled transect |
| 34 | b.tran.q095_green | transect | green channel pixel intensity at the 95% on the scaled transect |
| 35 | b.mean_red | intensity | red channel mean intensity |
| 36 | b.mean_blue | intensity | red channel standard deviation intensity |
| 37 | b.sd_red | intensity | red channel mad intensity |
| 38 | b.sd_blue | intensity | blue channel mean intensity |
| 39 | b.mad_red | intensity | blue channel standard deviation intensity |
| 40 | b.mad_blue | intensity | blue channel mad intensity |

**Supplementary Table 2. Isolates and image file names for monoculture and colony counting.** The column "Time" represents the transfer when the monoculture images were taken.

| Batch | Community | Isolate | Time | Image name | Colony count |
|---|---|---|---|---|---|
| B2 | C8R4 | 1 | T8 | B2_T8_C8R4_1 | 146 |
| B2 | C8R4 | 2 | T8 | B2_T8_C8R4_2 | 43 |
| B2 | C8R4 | 3 | T8 | B2_T8_C8R4_3 | 208 |
| B2 | C10R2 | 1 | T8 | B2_T8_C10R2_1 | 34 |
| B2 | C10R2 | 2 | T8 | B2_T8_C10R2_2 | 106 |
| B2 | C10R2 | 3 | T0 | B2_T0_C10R2_3 | 356 |
| B2 | C2R6 | 1 | T1 | B2_T1_C2R6_1 | 85 |
| B2 | C2R6 | 2 | T8 | B2_T8_C2R6_2 | 48 |
| B2 | C2R6 | 3 | T8 | B2_T8_C2R6_3 | 132 |
| B2 | C2R6 | 4 | T8 | B2_T8_C2R6_4 | 24 |
| B2 | C2R8 | 1 | T8 | B2_T8_C2R8_1 | 50 |
| B2 | C2R8 | 2 | T8 | B2_T8_C2R8_2 | 72 |
| B2 | C2R8 | 3 | T0 | B2_T0_C2R8_3 | 161 |
| B2 | C2R8 | 4 | T0 | B2_T0_C2R8_4 | 279 |
| B2 | C7R1 | 1 | T8 | B2_T8_C7R1_1 | 66 |
| B2 | C7R1 | 2 | T8 | B2_T8_C7R1_2 | 27 |
| B2 | C7R1 | 3 | T8 | B2_T8_C7R1_3 | 62 |
| B2 | C7R1 | 4 | T8 | B2_T8_C7R1_4 | 60 |
| B2 | C11R1 | 2 | T0 | B2_T0_C11R1_2 | 1,049 |
| B2 | C11R1 | 3 | T1 | B2_T1_C11R1_3 | 8 |
| B2 | C11R1 | 4 | T1 | B2_T1_C11R1_4 | 27 |
| B2 | C11R1 | 5 | T8 | B2_T8_C11R1_5 | 73 |
| B2 | C11R1 | 6 | T8 | B2_T8_C11R1_6 | 70 |
| B2 | C11R1 | 7 | T8 | B2_T8_C11R1_7 | 119 |
| B2 | C11R1 | 8 | T1 | B2_T1_C11R1_8 | 3 |
| B2 | C11R1 | 9 | T0 | B2_T0_C11R1_9 | 195 |
| C | C11R1 | 1 | T0 | C_T0_C11R1_1 | 8 |
| C2 | C11R2 | 1 | T8 | C2_T8_C11R2_1 | 54 |
| C2 | C11R2 | 2 | T0 | C2_T0_C11R2_2 | 80 |
| C2 | C11R2 | 3 | T8 | C2_T8_C11R2_3 | 42 |
| C2 | C11R2 | 4 | T8 | C2_T8_C11R2_4 | 120 |
| C2 | C11R2 | 5 | T8 | C2_T8_C11R2_5 | 62 |
| C2 | C11R2 | 6 | T8 | C2_T8_C11R2_6 | 128 |
| C2 | C11R2 | 7 | T8 | C2_T8_C11R2_7 | 28 |
| C2 | C11R2 | 8 | T0 | C2_T0_C11R2_8 | 85 |

| Batch | Community | Isolate | Time | Image name | Colony count |
|---|---|---|---|---|---|
| C2 | C11R2 | 9 | T0 | C2_T0_C11R2_9 | 58 |
| C2 | C11R2 | 10 | T0 | C2_T0_C11R2_10 | 70 |
| C2 | C11R2 | 11 | T8 | C2_T8_C11R2_11 | 66 |
| C2 | C11R2 | 12 | T8 | C2_T8_C11R2_12 | 37 |
| D | C4R1 | 1 | T8 | D_T8_C4R1_1 | 47 |
| D | C4R1 | 2 | T8 | D_T8_C4R1_2 | 146 |
| D | C4R1 | 3 | T8 | D_T8_C4R1_3 | 27 |
| D | C1R2 | 1 | T8 | D_T8_C1R2_1 | 82 |
| D | C1R2 | 2 | T8 | D_T8_C1R2_2 | 92 |
| D | C1R2 | 3 | T8 | D_T8_C1R2_3 | 130 |
| D | C1R2 | 4 | T8 | D_T8_C1R2_4 | 172 |
| D | C1R4 | 1 | T8 | D_T8_C1R4_1 | 62 |
| D | C1R4 | 2 | T8 | D_T8_C1R4_2 | 103 |
| D | C1R4 | 3 | T0 | D_T0_C1R4_3 | 75 |
| D | C1R4 | 4 | T8 | D_T8_C1R4_4 | 22 |
| D | C1R4 | 5 | T8 | D_T8_C1R4_5 | 41 |
| D | C1R6 | 1 | T8 | D_T8_C1R6_1 | 46 |
| D | C1R6 | 2 | T8 | D_T8_C1R6_2 | 105 |
| D | C1R6 | 3 | T0 | D_T0_C1R6_3 | 31 |
| D | C1R6 | 4 | T8 | D_T8_C1R6_4 | 109 |
| D | C1R6 | 5 | T8 | D_T8_C1R6_5 | 69 |
| D | C11R5 | 1 | T8 | D_T8_C11R5_1 | 38 |
| D | C11R5 | 2 | T8 | D_T8_C11R5_2 | 123 |
| D | C11R5 | 3 | T8 | D_T8_C11R5_3 | 58 |
| D | C11R5 | 4 | T8 | D_T8_C11R5_4 | 111 |
| D | C11R5 | 5 | T8 | D_T8_C11R5_5 | 4 |
| D | C1R7 | 1 | T8 | D_T8_C1R7_1 | 135 |
| D | C1R7 | 2 | T8 | D_T8_C1R7_2 | 40 |
| D | C1R7 | 3 | T8 | D_T8_C1R7_3 | 35 |
| D | C1R7 | 4 | T8 | D_T8_C1R7_4 | 168 |
| D | C1R7 | 5 | T8 | D_T8_C1R7_5 | 54 |
| D | C1R7 | 6 | T8 | D_T8_C1R7_6 | 46 |
| D | C1R7 | 7 | T1 | D_T1_C1R7_7 | 90 |

**Supplementary Table 3. Pairwise competition outcome with frequency changes.** The minus sign (-) means a statistically significant decrease from the initial frequency to the final frequency, the plus sign (+) indicates a statistically significant increase and a zero (0) means no statistical significance in the change in frequency.

| | From rare | From medium | From abundant | Outcome | Finer outcome | Count |
|---|---|---|---|---|---|---|
| 1 | + | + | + | exclusion | competitive exclusion | 48 |
| 2 | + | + | - | coexistence | stable coexistence | 10 |
| 3 | + | + | 0 | coexistence | coexistence at 95% | 20 |
| 4 | + | - | + | coexistence | frequency-dependent coexistence | 0 |
| 5 | + | - | - | coexistence | stable coexistence | 14 |
| 6 | + | - | 0 | coexistence | frequency-dependent coexistence | 1 |
| 7 | + | 0 | + | unknown | | 3 |
| 8 | + | 0 | - | coexistence | stable coexistence | 8 |
| 9 | + | 0 | 0 | coexistence | 2-freq neutrality | 2 |
| 10 | - | + | + | exclusion | mutual exclusion | 0 |
| 11 | - | + | - | coexistence | frequency-dependent coexistence | 1 |
| 12 | - | + | 0 | unknown | | 0 |
| 13 | - | - | + | exclusion | mutual exclusion | 0 |
| 14 | - | - | - | exclusion | competitive exclusion | 48 |
| 15 | - | - | 0 | unknown | | 3 |
| 16 | - | 0 | + | unknown | | 0 |
| 17 | - | 0 | - | unknown | | 2 |
| 18 | - | 0 | 0 | coexistence | 2-freq neutrality | 0 |
| 19 | 0 | + | + | unknown | | 0 |
| 20 | 0 | + | - | coexistence | frequency-dependent coexistence | 1 |
| 21 | 0 | + | 0 | coexistence | 2-freq neutrality | 0 |
| 22 | 0 | - | + | unknown | | 0 |
| 23 | 0 | - | - | coexistence | coexistence at 5% | 18 |
| 24 | 0 | - | 0 | coexistence | 2-freq neutrality | 0 |
| 25 | 0 | 0 | + | coexistence | 2-freq neutrality | 0 |
| 26 | 0 | 0 | - | coexistence | 2-freq neutrality | 1 |
| 27 | 0 | 0 | 0 | coexistence | 3-freq neutrality | 1 |
